# Conservation and diversity in transcriptional responses among host plants forming distinct arbuscular mycorrhizal morphotypes

**DOI:** 10.1101/2021.06.07.447186

**Authors:** Takaya Tominaga, Chihiro Miura, Yuuka Sumigawa, Yukine Hirose, Katsushi Yamaguchi, Shuji Shigenobu, Akira Mine, Hironori Kaminaka

## Abstract

- The morphotype of arbuscular mycorrhizal (AM) roots is distinct mostly depending on AM host species: *Arum*, *Paris*, and Intermediate types. We previously reported that gibberellin (GA) promotes the establishment of *Paris*-type AM symbiosis in *Eustoma grandiflorum* despite its negative effects on *Arum*-type AM symbiosis in model plants. However, the molecular mechanisms underlying the differential effects of GA on different morphotypes, including Intermediate-type AM symbiosis, remain elusive.
- Comparative transcriptomics revealed that several symbiosis-related genes were transcriptionally promoted upon AM fungal colonization in *Lotus japonicus* (*Arum*-type), *Daucus carota* (Intermediate-type), and *E. grandiflorum* (*Paris*-type). Interestingly, upon GA treatment, the fungal colonization levels and expression of symbiosis-related genes were suppressed in *L. japonicus* and *D. carota* but were promoted in *E. grandiflorum*.
- Exogenous GA transcriptionally inhibited the biosynthetic process of a host-derived signal molecule involved in AM symbiosis, strigolactone, in *L. japonicus* and *E. grandiflorum*. Additionally, disaccharides mainly metabolized in AM roots would be different between *L. japonicus* and *D. carota*/*E. grandiflorum*.
- This study uncovered the conserved transcriptional responses during mycorrhization and diverse responses to GA in AM roots with distinct morphotypes among phylogenetically distant host plants.

## Introduction

More than 70% of terrestrial plants associate with the symbiotic fungi, arbuscular mycorrhizal (AM) fungi, belonging to Glomeromycotina (Brundrett & Tedersoo, 2018). AM fungi supply minerals, such as inorganic phosphate and nitrogen, to their host plants, thus promoting the growth of the hosts (Ezawa & Saito, 2018; Wang *et al*., 2020). In return, as AM fungi cannot produce carbohydrates, they obtain carbohydrates, such as fatty acids, lipids, and monosaccharides, from the host plants (Bravo *et al*., 2017; Kobayashi *et al*., 2018; An *et al*., 2019). This mutual interaction is established through several steps. Before establishing AM symbiosis, the host-derived signal molecules, strigolactones (SLs), exudate into the rhizosphere to attract AM fungi (Akiyama *et al*., 2005; Besserer *et al*., 2008; Tsuzuki *et al*., 2016). SLs are also required to continuously form hyphopodia on the host root epidermis (Kobae *et al*., 2018). After AM fungal hyphae invade the host epidermis, AM fungi form highly branched hyphal structures, the arbuscule, in the root cortical cells for nutrient exchange. Some transporters are localized on a specialized plant-derived membrane, periarbuscular membrane (PAM), to influx mineral nutrients and efflux carbohydrates, such as lipids and glucose, between fungal symbionts (Kobae & Hata, 2010; Bravo *et al*., 2017; Luginbuehl & Oldroyd, 2017).

AM symbiosis has been known to be altered by the phytohormone, gibberellin (GA). Exogenous GA severely reduces the number of hyphopodia and disturbs the development of arbuscule (Floss *et al*., 2013; Yu *et al*., 2014; Takeda *et al*., 2015; Pimprikar *et al*., 2016). Moreover, GA represses the expressions of some AM symbiosis-related genes (Takeda *et al*., 2015; Pimprikar *et al*., 2016). Notably, it has shown that a GRAS transcription factor (TF) required for AM symbiosis, *REDUCED ARBUSCULAR MYCORRHIZA1* (*RAM1*), is transcriptionally downregulated in GA-treated *Lotus japonicus*. This is attributable to the GA-triggered degradation of GA signaling repressor, DELLA, positively regulating *RAM1* expression (Silverstone *et al*., 2001; Achard & Genschik, 2009; Floss *et al*., 2013; Park *et al*., 2015). Notably, the RAM1 also regulates other downstream AM marker genes: *REDUCED FOR ARBUSCULE DEVELOPMENT1* (*RAD1*)–GRAS TF, *Vapyrin* (*Vpy*) (protein that regulates arbuscule development), *PHOSPHATE TRANSPORTER4* (*PT4*), *AMMONIUM TRANSPORTER2;2* (*AMT2;2*), *FatM* (acyl-acyl carrier protein thioesterase), *RAM2* (glycerol-3-phosphate acyltransferase), and *STR/STR2* (ABC transporters for lipids) (Gobbato *et al*., 2013; Park *et al*., 2015; Rich *et al*., 2015; Pimprikar *et al*., 2016; Rich *et al*., 2017; Muller *et al*., 2020). Thereby, it has been thought that exogenous GA or the absence of functional DELLA attenuates the transcriptional promotion of downstream genes, inhibiting AM fungal colonization.

In contrast to the negative effects of GA on *Arum*-type AM symbiosis in *L. japonicus* and chive, we previously reported the positive effects of GA treatment on *Paris*-type AM symbiosis in *Eustoma grandiflorum* and *Primula malacoides* (Tominaga *et al*., 2020a). Previous expression analysis also showed that the expression levels of AM symbiosis-related genes in *E. grandiflorum* were promoted by GA (Tominaga *et al*., 2020b). *Arum*-type AM shows that AM fungal hyphae elongate intercellular space of the host cortex and form arbuscules in the cortical cells. This AM morphotype is found in rice (*Oryza sativa*) and legume model plants roots, and generally seen in recent reports (Yu *et al*., 2014; Takeda *et al*., 2015). On the other hand, in *Paris*-type AM, the fungal hyphae invade through and show arbuscule-branched hyphal coils in the cortical cells (Smith & Smith, 1997; Dickson, 2004; Dickson *et al*., 2007). However, our previous studies did not simultaneously compare the GA-mediated transcriptional regulation between *E. grandiflorum* and other host plants at the molecular level. Moreover, some host plants show an “Intermediate” type of AM; linear AM fungal hyphae intracellularly elongate (Dickson, 2004). To date, the effect of GA on Intermediate-type AM symbiosis has not been investigated.

In this study, we conducted comparative transcriptomics among three AM host plants with different AM morphotypes: *L. japonicus* (*Arum*-type AM), *E. grandiflorum* (*Paris*-type AM), and *Daucus carota* (Intermediate-type AM) (Dickson, 2004). Based on plastid genomes, Fabales, Gentianales, and Apiales, to which *L. japonicus*, *D. carota*, and *E. grandiflorum* belong, are estimated to appear *c*. 100 Ma, *c*. 90 Ma, and *c*. 80 Ma, respectively (Li *et al*., 2019). Our study revealed that *Rhizophagus irregularis* infection promoted shoot growth and the expression of several symbiosis-related genes conserved in all examined plants. However, the AM fungus-promoted expressions of the orthologs were decreased in *L. japonicus* (*Arum-*type) and *D. carota* (Intermediate-type) but further increased in *E. grandiflorum* (*Paris*-type) by exogenous GA. Additionally, the negative effects of GA on SL biosynthetic process were commonly observed in *L. japonicus* and *E. grandiflorum*, indicating that SLs would not be involved in GA-promoted fungal colonization in GA-treated *E. grandiflorum*. Therefore, our study uncovered the conserved responses during AM symbiosis regardless of AM morphotypes. Furthermore, our findings help understand that GA would be a key regulator showing diverse effects on AM symbioses depending on the host species as AM morphotypes demonstrate.

## Materials and Methods

### The growth condition of plant and fungal materials

The seedlings of *L. japonicus* “Miyakojima” MG-20, *D. carota* cv. Nantes, and *E. grandiflorum* cv. Pink Thumb were prepared as in our previous report (Tominaga *et al*., 2020a). *Daucus carota* seedlings were grown in light for 7 d. For colonization tests, approximately 6000 spores of *R. irregularis* DAOM197198 (Premier Tech, Quebec, Canada) were added to 50 ml 1/5 Hoagland solution containing 20 µM inorganic phosphate. GA_3_ was dissolved in ethanol and treated at this procedure by diluting the stock to the 1/5 Hoagland solution at indicated concentrations. Ethanol was treated in the same way as the control treatment. The solution was added to approximately 300 ml autoclaved mixed soil (river sand/vermiculite, 1:1) in a plastic container combined with another one as described in Takeda *et al*. (2015). Then, the prepared seedlings were transplanted into the soil and kept for 6 wk under 14 h light/10 h dark cycles at 25°C.

### Quantification and observation of AM symbiosis

The inoculated roots were harvested at 6 wk post inoculation (wpi), and fixation, staining, and quantification of AM fungal colonization rates were conducted according to previous studies (Mcgonigle *et al*., 1990; Tominaga *et al*., 2020a).

For fluorescence images, the fragments of fixed roots were rinsed with phosphate-buffered saline (PBS) and immersed in ClearSee (FUJIFLIM Wako Pure Chemical, Osaka, Japan) for 1 wk in the dark (Kurihara *et al*., 2015). The manufacturer’s instructions were followed in the clearing procedure. The cleared roots were rinsed with PBS and stained with 0.01 mg ml^−1^ WGA-Alexa Fluor 488 (Thermo Fisher Scientific, MA, USA) for 15 min. Under a fluorescent stereomicroscope, Leica M205 FCA (Leica Microsystems, Wetzlar, Germany), the relatively bright green fluorescent region, which indicates colonized area, was excised with a scalpel and embedded in 5% (w/v) agarose containing 1% (w/v) gelatin. Then, 30–50 µm cross sections were made using a Linear Slicer PRO-7 (Dosaka EM, Kyoto, Japan) and observed under a FLUOVIEW FV10i confocal laser scanning microscope (Olympus, Tokyo, Japan) with 499 nm excitation and 520 nm emission for WGA-Alexa Fluor 488 and FV10i-SW software v1.2 (Olympus). The images were merged using the ImageJ software v1.51k (http://imagej.nih.gov/ij).

### Transcriptome analysis

#### RNA extraction and RNA-seq

Sample roots (approximately 100 mg) at 6 wpi were collected in a nuclease-free tube (INA-OPTIKA, Osaka, Japan) with two 5 mm beads, frozen by liquid nitrogen. The frozen root samples were set in ShakeMan6 (Bio-Medical Science, Tokyo, Japan) and homogenized. Then, the total RNA was extracted using the real RNA Extraction Kit Mini for Plants (RBC Bioscience, New Taipei, Taiwan) following the manufacturer’s protocol. RNase-free DNase I (Takara Bio, Shiga, Japan) was applied to remove genomic DNA from the RNA samples according to the manufacturer’s instructions. The purity and quantity of the total RNA was calculated by measuring the absorbance at 260 and 280 nm (*A*260: *A*280) with DeNovix DS-11 + (Scrum, Tokyo, Japan). RNA-seq library was constructed from the total extracted RNA and sequenced, and RNA-seq with strand-specific and paired-end reads (150 bp) was performed with DNBSEQ-G400 by Genewiz (Tokyo, Japan). Consequently, more than 20 million raw reads per sample were obtained (Table S1). Low-quality reads (< QV30) and adapter sequences were removed by Fastp (Chen *et al*., 2018).

#### Data analysis

Read mapping was conducted using STAR (Dobin *et al*., 2013) for the filtered single-end reads of *L. japonicus*, *D. carota*, and *R. irregularis* onto their genomes, Lotus japonicus Lj1.0v1, Daucus carota v2.0, and Rir_HGAP_ii_V2, retrieved from the Phytozome v13 (https://phytozome-next.jgi.doe.gov) and Ensembl Fungi (Iorizzo *et al*., 2016; Maeda *et al*., 2018; Li *et al*., 2020). Meanwhile, Bowtie2 with default parameters except for “--local” was applied for *E. grandiflorum* to map the reads to *de novo* reference assembly constructed from previous RNA-seq data (Tominaga *et al*., 2020b) by Trinity v2.8.4 (Grabherr *et al*., 2011; Langmead & Salzberg, 2012; Haas *et al*., 2013). In this study, we mapped the reverse reads to the indicated genomes or *de novo* assembly data to perform specific alignment. The number of mapped reads to the reference genome was counted using featureCounts v1.6.4 for *L. japonicus*, *D. carota*, and *R. irregularis*, whereas that of *E. grandiflorum* was quantified with eXpress v1.5.1 (Roberts & Pachter, 2013; Liao *et al*., 2014). For identifying differentially expressed genes (DEGs), each count data showing different library sizes was normalized by the trimmed mean of the *M*-values normalization method, and genes with |Log_2_ fold change (FC)| > 1 and FDR less than the indicated values (FDR < 0.01 for plants’ DEGs and FDR < 0.05 for fungal DEGs) were considered DEGs using the EdgeR package (Robinson *et al*., 2010).

The transcripts per million (TPM) (Li *et al*., 2010; Wagner *et al*., 2012) of each sample was counted from the count data using the R software v4.0.2 (R Foundation for Statistical Computing). Genes that showed zero counts in all samples were excluded from the analysis, unless otherwise noted. Then, the mean TPM of all samples in a condition was Log_2_-transformed for each gene. The heatmaps in this study were constructed using the heatmaply package in R (Galili *et al*., 2018).

#### Gene ontology enrichment analysis

DEG was sorted depending on their expression patterns using a Venn diagram (http://bioinformatics.psb.ugent.be/webtools/Venn). Then, gene ontology (GO) enrichment analysis was conducted using the ClueGO plugin for Cytoscape (Bindea *et al*., 2009; Bindea *et al*., 2013). Additionally, the correlation network of enriched GO terms was created using the ClueGO. In the analysis, *P*-values were calculated using a two-sided hypergeometric test and corrected using the Benjamini–Hochberg method. The GO terms of *R. irregularis* were annotated by EnTAP v0.10.7 (Hart *et al*., 2020), followed by GO enrichment analysis using the topGO package in the R environment. In the topGO study, the enrichment test was performed by calculating the *P*-values using the Kolmogorov–Smirnov test (*P <* 0.01) and scoring with the *Elim* method (Alexa *et al*., 2006).

### Ortholog identification

Here, we identified ortholog genes in *L. japonicus*, *D. carota*, and *E. grandiflorum* to compare the influence of AM fungal colonization and GA treatment among these different host species. The proteomes of *L. japonicus* and *D. carota* was retrieved from the Phytozome v12.1 and v13, respectively. Additionally, coding sequence and amino acid sequences in the *de novo* assembly of *E. grandiflorum* were predicted using TransDecoder v5.5.0 (Haas *et al*., 2013). Next, the ortholog was identified using SonicParanoid with default parameters in the Python v3.8 environment (Cosentino & Iwasaki, 2019). Several known genes were used as queries for BLASTp search against *L. japonicus* proteome on the website Phytozome v13 (Tables S2, S3). The resulting top hit *L. japonicus* gene and its corresponding orthologs in *D. carota* and *E. grandiflorum* were considered orthogroups and analyzed.

### Extraction of endogenous SLs and germination assay

To extract SLs from the host roots, we referred to the methods in a previous study with some modifications (Floková *et al*., 2020). The fresh 6-wk-old roots (100 mg) were homogenized in ShakeMan6 (Bio-Medical Science) with 1 ml of 60% (v/v) acetone stored at −30°C. The suspensions were collected by centrifugation and evaporated *in vacuo* for 30 min using Savant SpeedVac DNA130 (Thermo Fisher). Hydrophobic components in residual water (*c*. 500 µl) were extracted by ethyl acetate three times, and the organic layer was evaporated *in vacuo*. The samples were resolved in acetone at 400 mg FW root ml^-1^ and stored at 4°C until use.

*Orobanche minor* seeds were incubated on two moist filter papers for 10 d at 24°C in the dark. An aliquot of acetone, 1 µM *rac*-GR24 (StrigoLab, Torino, Italy), and extracted samples (20 µl) were added to 6 mm glass fiber disks. Then, the conditioned *O. minor* seeds were placed on the disks with 20 µl distilled water. After 5 d of incubation at 24°C in the dark, the germination rate (%) was counted.

### Biological replicate and statistical analysis

One glass slide with 10 pieces of root fragments collected from one plant was considered a biological replicate for colonization rate quantification. One glass fiber disc with *O. minor* seeds was equivalent to one biological replicate. These experiments were reproduced three times with more than five biological replicates. In the transcriptome analysis, one library constructed from a pool of total RNA consisting of three plants was treated as one biological replicates. Statistical analyses were conducted using the R software v4.0.2.

## Results

### Phenotypes of arbuscular mycorrhizal roots in different host plant species associated with *Rhizophagus irregularis*

In *L. japonicus*, a typical *Arum*-type AM with intercellular hyphae and highly branched arbuscules in the cortical cells were formed at 6 wpi (Fig. 1a). We also observed *D. carota* AM roots and found linear intraradical hyphae invading the cortical cells, but no hyphal coil was formed (Fig. 1c). These morphological features in *D. carota* roots are Intermediate-type AM in association with another AM fungus, *Glomus mosseae* (Dickson, 2004). Additionally, *E. grandiflorum* showed a classical *Paris*-type AM that forms hyphal coils with arbuscule branching and intracellular hyphae penetrating the cortical cells (Fig. 1e) (Tominaga *et al*., 2020a). Therefore, our result confirmed that the associating plant species mainly determine the AM morphologies in the same monoxenic culture conditions, although some exceptions are reported (Cavagnaro *et al*., 2001; Dickson, 2004; Smith *et al*., 2004; Kubota *et al*., 2005; Hong *et al*., 2012).

**Fig. 1.**
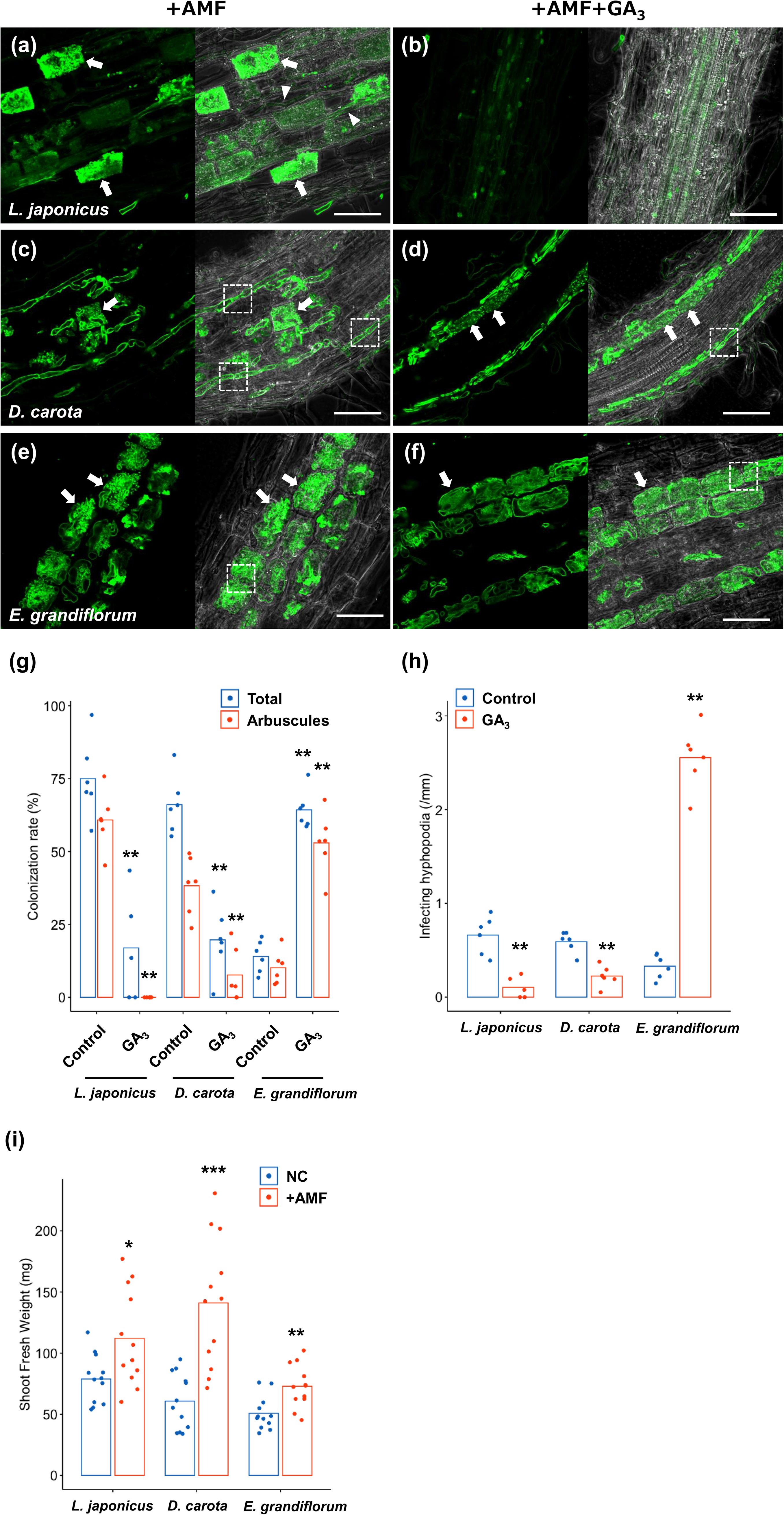
Observation of AM morphotypes and quantification of AM fungal colonization levels and plant growth promotion. AM symbiosis-related phenotypes and responses to GA in the examined host plants colonized by *R. irregularis* were observed at 6 wpi. Each host plant was treated with 0.01% ethanol (+AMF) for the control and 1 µM GA_3_ (+AMF+GA_3_). The collected AM roots were stained with WAG-Alexa Fluor 488 (green). (a–f) Confocal images of *L. japonicus* (a, b), *D. carota* (c, d), and *E. grandiflorum* (e, f) inoculated with *R. irregularis* in the +AMF (a, c, e) and +AMF+GA_3_ condition (b, d, f). The left sides of each confocal fluorescence image are merged with their images in bright field (right sides). Arrows, arbuscules; arrowheads, intercellular hyphae; dotted lines, intracellular hyphae penetrating two adjacent cortical cells; bars, 200 µm in (b) and 50 µm in the others. (g) Quantification of AM fungal colonization rates (%) in 0.01% ethanol- and GA-treated roots of the examined plants. Total, total colonization rate; arbuscules, percentage of arbuscule formation (n = 5, 6). (h) The number of hyphopodia penetrating the host root epidermis per root length (mm) (n = 5, 6). (i) Shoot fresh weight (mg) of each host plant grown in axenic conditions (NC) and colonized by *R. irregularis* (+AMF) (n = 12). The average percentage values are shown as bars. Asterisks indicate significant differences compared with the controls in the Wilcoxon rank-sum test (*: *P <* 0.05, **: *P <* 0.01, ***: *P <* 0.001).

Although it has been reported that exogenous GA treatment differentially alters the establishment of *Arum*- and *Paris*-type AM symbiosis (Tominaga *et al*., 2020a), how *D. carota*–*R. irregularis* interaction responses to GA has not been tested yet. We treated *L. japonicus*, *D. carota*, and *E. grandiflorum* with 1 µM GA_3_ and quantified fungal colonization. The rates of total hyphal structures and arbuscules were significantly decreased in GA-treated *L. japonicus* but increased in GA-treated *E. grandiflorum* compared with the control roots (Fig. 1g). In GA-treated *L. japonicus* roots, we could not find any arbuscules, but we found some intercellular hyphae as described in several studies (Fig. 1b, g) (Takeda *et al*., 2015; Pimprikar *et al*., 2016). Interestingly, the morphologies of hyphal structures in *D. carota* and *E. grandiflorum* were not influenced by GA treatment (Fig. 1d, f). These results indicate that the two plants can form normal arbuscules, even in the presence of GA, implying that *L. japonicus* was more sensitive to GA than *D. carota*. The number of AM fungal entries was consistent with the colonization rates (Fig. 1h). Additionally, AM fungal colonization commonly promoted the growth of examined host plants in the control condition (Fig. 1i). Therefore, associating with *R. irregularis* contributed to growth promotion in each tested plant regardless of AM morphotypes, whereas the responses to exogenous GA in *E. grandiflorum* AM roots were unique.

### Comparisons of symbiosis-related genes shed light on conserved and specific transcriptional responses among AM host plants

Based on Fig. 1, the transcriptional regulation of downstream genes required for AM symbiosis would be different among the examined plants (Fig. 1g). To test this hypothesis, the expression pattern of genes conserved among the host plants was compared. First, orthogroups, including each known AM symbiosis-related gene, were identified using the SonicParanoid software (Tables S2, S3).

We focused on several downstream genes in AM symbiosis: *RAM1*, *RAD1*, *Vpy*, *PT4*, *AMT2;2, AMT2;3*, *FatM*, *RAM2*, *STR*, and *STR2* (Fig. 2a; Tables S2, S3). *CBX1* and *WRI5a/b/c*, AP2/EREBP domain TFs that positively regulate *RAM1* expression and fatty acid biosynthesis, were also included in the analysis (Luginbuehl *et al*., 2017; Jiang *et al*., 2018; Xue *et al*., 2018). Additionally, we identified the sucrose synthase 1 (*SucS1*) and glucose transporter (Sugar Will Eventually be Exported Transporter 1b; *SWEET1b*) conserved in the examined plants. *SucS1* and *SWEET1b* are predicted to catalyze sucrose into glucose in arbuscule-containing cortical cells and export the monosaccharide on PAM in *Medicago truncatula*, respectively (Hohnjec *et al*., 2003; Baier *et al*., 2010; An *et al*., 2019). Notably, these genes have been reported to be transcribed upon AM fungal colonization.

**Fig. 2.**
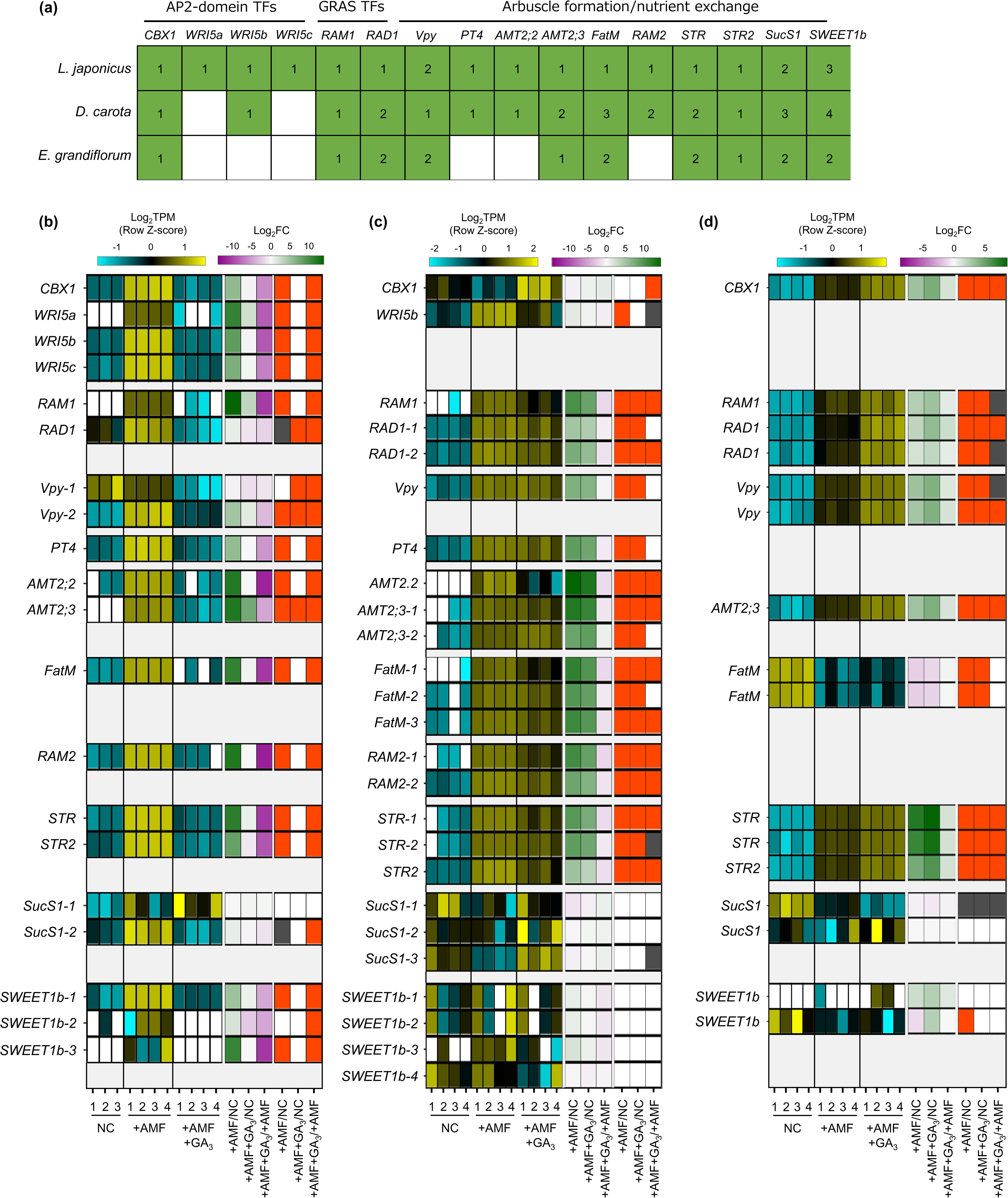
Comparative analysis on the expression patterns of AM symbiosis-related downstream genes conserved among examined host plants. (a) A table showing AM symbiosis-related downstream genes in *L. japonicus*, *D. carota*, and *E. grandiflorum* identified using the SonicParanoid. Green represents the conservation of respective gene, and the values show the number of homologous genes and transcripts in *L. japonicus*/*D. carota* genomes and the *de novo* assemble data of *E. grandiflorum*, respectively (Table S3). (b–d) Heatmaps represent the expression patterns of the selected genes in response to AM fungal colonization and GA treatment in *L. japonicus* (b), *D. carota* (c), and *E. grandiflorum* (d) at 6 wpi. The left heatmaps indicate the expression levels of selected genes. The Log-transformed TPM in every sample is shown in blue (low expression level), black (mean), and yellow (high expression level); the Z-score is normalized to turn the average value and SD to 0 and 1, respectively, across all samples. The number below the heatmaps indicates biological replicate. The middle ones show Log_2_-transformed fold changes in the genes compared with the controls. Magenta indicates negative values, green represents positive values, and white means 0. The right ones illustrate significance in the fold changes in gene expression levels. DEGs (|Log_2_FC| > 1, FDR < 0.01) and genes showing significantly but slightly different expression levels compared with the controls (FDR < 0.05) are colored with red and grey, respectively. NC, non-colonized control; +AMF, *R. irregularis* inoculation; +AMF+GA_3_, simultaneous application of *R. irregularis* inoculation and 1 µM GA_3_. The DEGs were identified by comparing +AMF with NC (+AMF/NC), +AMF+GA_3_ against NC or +AMF (+AMF+GA_3_/NC, +AMF+GA_3_/+AMF). For the TPM, Log_2_FC, and FDR values of the selected gene, see Table S4.

In this analysis, the examined plants were grown under several conditions as follows: non-colonized control roots (NC), AM roots (+AMF), and GA-treated AM roots (+AMF+GA_3_). A common set of selected genes were transcriptionally promoted upon fungal colonization at 6 wpi in each plant (Fig. 2b–d). AM fungal colonization, however, did not induce the expression of *FatM*, *SucS1*, and *SWEET1b* in *D. carota* or *E. grandiflorum* or both at the time point (Fig. 2c, d). In *L. japonicus*, the expression levels of several conserved genes were undetectable or mostly reduced by exogenous GA compared with NC and +AMF conditions (Fig. 2b). In contrast, the expression levels of AM symbiosis-related genes in GA-treated *D. carota* were still increased compared with the NC but decreased compared with the +AMF (Fig. 2c). This indicates that the negative effect of GA on *D. carota* would be relatively moderate to that on *L. japonicus* as the colonization rates suggested (Fig. 1g). As for *E. grandiflorum*, the expression of the AM-induced genes was further enhanced by GA than the NC and +AMF controls (Fig. 2d). This result also supports the quantitative analysis of fungal colonization (Fig. 1g).

In this study, the SonicParanoid could not find *E. grandiflorum PT4* (*EgPT4*) and *RAM2* (*EgRAM2*). Therefore, we also conducted a BLAST search for the two genes in *E. grandiflorum* with sufficient E-value (<1E-5) (Table S4). Consequently, one gene annotated as phosphate transporter (TRINITY_DN34977_c0_g1_i1.p1) was transcriptionally enhanced upon AM fungal colonization and exogenous GA (Table S4) (Tominaga *et al*., 2020b). However, representative genes described as glycerol-3-phosphate acyltransferase (*RAM2*) were not transcriptionally promoted upon AM fungal colonization (Table S4). Alternatively, we might have missed *EgRAM2* in the *de novo* assembly after removing redundant contigs with CD-HIT (Tominaga *et al*., 2020b).

### Genes involved in phytohormone biosynthesis and signaling show similar transcriptional responses to exogenous gibberellin in examined host plants

Since the number of infecting hyphopodia differed between *L. japonicus*/*D. carota* and *E. grandiflorum*, we also analyzed the expression patterns of several SL-related genes. *D27*, *CCD7*, *CCD8*, and *MAX1* are necessary for SL biosynthesis (Booker *et al*., 2004; Booker *et al*., 2005; Auldridge *et al*., 2006; Alder *et al*., 2012; Waters *et al*., 2012a; Al-Babili & Bouwmeester, 2015). *PDR1* in *Petunia hybrida* encoding a G-type ABC transporter is predicted to export SLs (Kretzschmar *et al*., 2012). Additionally, *D14*, *DLK2*, and *KAI2*, which belong to a D14 family, have previously been demonstrated as components in SL or karrikin signaling or both (Waters *et al*., 2012b; Kameoka & Kyozuka, 2015; Vegh *et al*., 2017). Based on the identification using SonicParanoid, these genes seemed to be conserved in all host plants (Fig. 3a; Table S3). For SL biosynthetic genes, *LjD27*, *LjCCD7*, and *LjCCD8* were transcriptionally downregulated by GA treatment (Fig. 3b). Additionally, the expression pattern of these genes in *E. grandiflorum* was similar to that in *L. japonicus*, suggesting that GA inhibits SL biosynthesis and exudation to the rhizosphere in *E. grandiflorum* as model plants (Fig. 3d) (Ito *et al*., 2017).

**Fig. 3.**
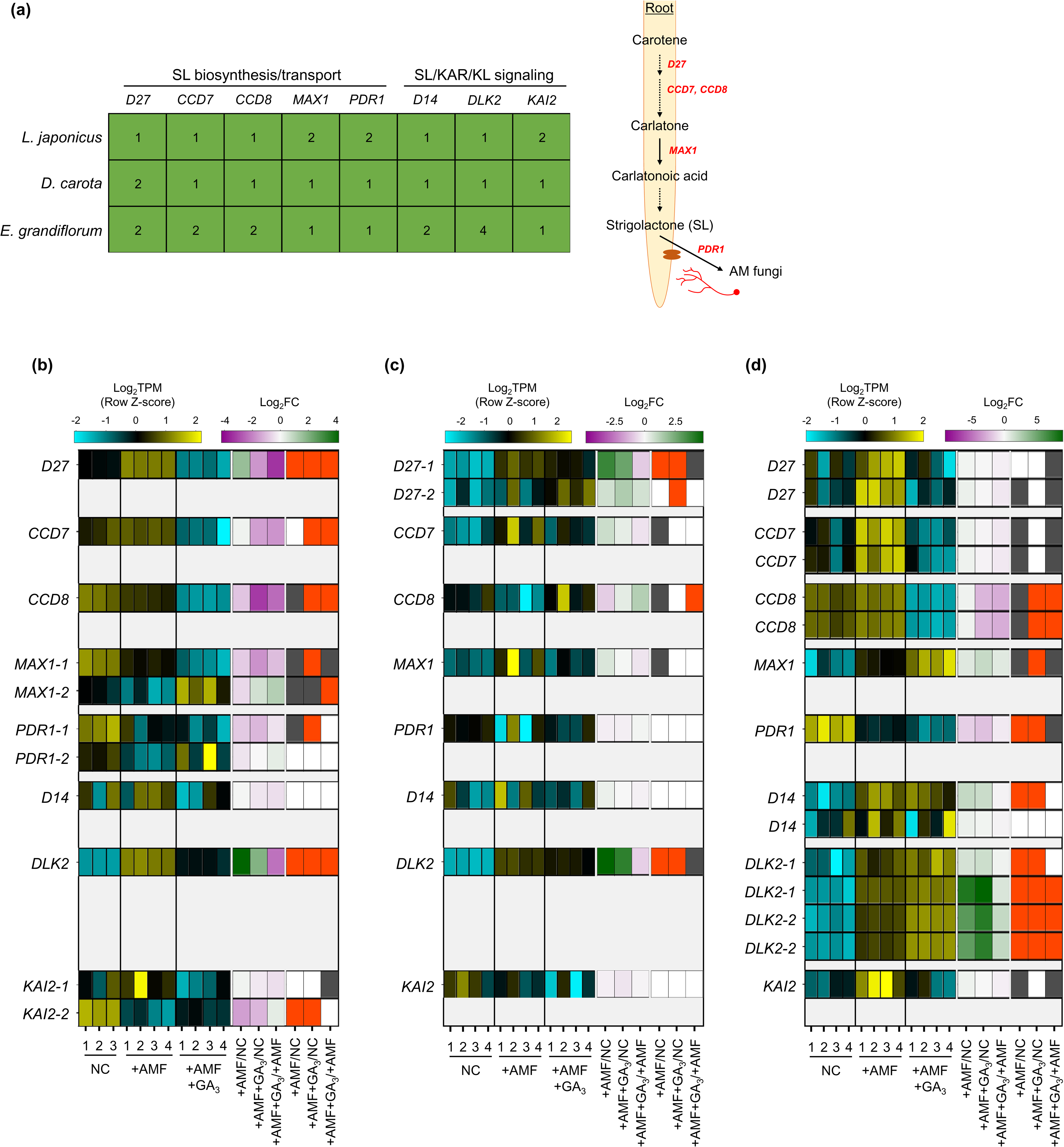
Comparative analysis of the expression patterns of SL biosynthetic and signaling-related genes conserved among examined host plants. (a) A table showing SL-related downstream genes in *L. japonicus* and *D. carota*, and transcripts in *E. grandiflorum* identified using the SonicParanoid as demonstrated in Fig. 2 (Output of SonicParanoid: Tables S2, S3). (a) This analysis was conducted at 6 wpi. The right image illustrates the biosynthetic process and exudation of SLs in the root. (b–d) The expression levels and Log_2_FC compared with the controls of each gene are represented in the left heatmap (blue, low expression level; black, mean; yellow, high expression level) and the middle one (magenta, negative values; green, positive values; white, zero), respectively. The number below the heatmaps indicates a biological replicate. In the right one, the red and grey represent DEGs (|Log_2_FC| > 1, FDR < 0.01) and genes with FDR < 0.05. NC, non-colonized control; +AMF, *R. irregularis* inoculation; +AMF+GA_3_, simultaneous application of *R. irregularis* inoculation and 1 µM GA_3_. The DEGs were identified by comparing +AMF with NC (+AMF/NC), +AMF+GA_3_ against NC or +AMF (+AMF+GA_3_/NC, +AMF+GA_3_/+AMF). For the TPM, Log_2_FC, and FDR values of the selected genes, see Table S4.

However, *DcCCD8* expression was significantly accelerated in GA-treated AM roots than the +AMF condition (Fig. 3c). To confirm the effect of GA on SL production, we conducted a germination assay by applying *O. minor* whose germination is induced by SLs (Ueno *et al*., 2014; Trabelsi *et al*., 2017). To prepare root extraction in the same conditions as the RNA-seq experiments, the examined plants were grown in the soil mixture for 6 wk. Consistent with the expression analysis, the germination activity of root extracts was significantly reduced but increased in *L. japonicus* and *D. carota* by GA treatment, respectively (Fig. S1). The seed germination of *O. minor* was not promoted by *E. grandiflorum* root extracts, which might be attributed to the low quantities of SLs (Sato *et al*., 2003; Halouzka *et al*., 2020). If we hydroponically grow *E. grandiflorum*, the root exudates exhibited germination activity (data not shown).

As for several SL signaling-related genes, AM fungal colonization significantly increased the expression of *DLK2* genes among all host plants compared with NC control (Fig. 3b–d). These results were similar in other host plants, *O. sativa* and *Solanum lycopersicum* (Choi *et al*., 2020; Ho-Plagaro *et al*., 2021). Moreover, compared with the +AMF control, GA treatment significantly reduced *DLK2* expression in *L. japonicus*; however, *DLK2* expression further increased in *E. grandiflorum* (Fig. 3b, d) (Tominaga *et al*., 2020b). The expression of *DLK2* in GA-treated *D. carota* was slightly reduced (Fig. 3c).

### Transcriptional responses to exogenous gibberellin reflect fungal colonization rates in three host plant species

For further effective comparison of transcriptional responses to AM fungal colonization and GA treatment, we also identified the ortholog genes with a one-to-one relationship among *L. japonicus*, *D. carota*, and *E. grandiflorum* using SonicParanoid. This resulted in 2705 ortholog genes (Table S2). Moreover, we selected DEGs that were expressed in all samples and met the criteria (|Log_2_FC| > 1 and FDR < 0.01) at least in one condition. In this analysis, the DEGs were identified by comparing +AMF with NC, +AMF+GA_3_ with NC, and +AMF+GA_3_ with +AMF conditions. Consequently, 221, 199, and 225 DEGs of the orthologs in *L. japonicus*, *D. carota*, and *E. grandiflorum* were arranged by mean Log_2_-transformed TPM as shown in a heatmap, respectively (Fig. 4a–c). In *L. japonicus* and *D. carota*, the expression pattern of the DEGs in +AMF+GA_3_ was grouped with NC but excluded from +AMF (Fig. 4a, b). However, the DEGs in +AMF+GA_3_ were grouped with +AMF in *E. grandiflorum* AM roots (Fig. 4c). Consistent with the results, *E. grandiflorum* in +AMF+GA_3_ condition was separated from *L. japonicus* and *D. carota* in the same condition when all 467 DEGs were arranged by their Log_2_FC (Fig. 4d). This hierarchical clustering by the Log_2_FC values further illustrated that *E. grandiflorum* in +AMF+GA_3_ condition and all plants in +AMF condition were included in the same cluster.

**Fig. 4.**
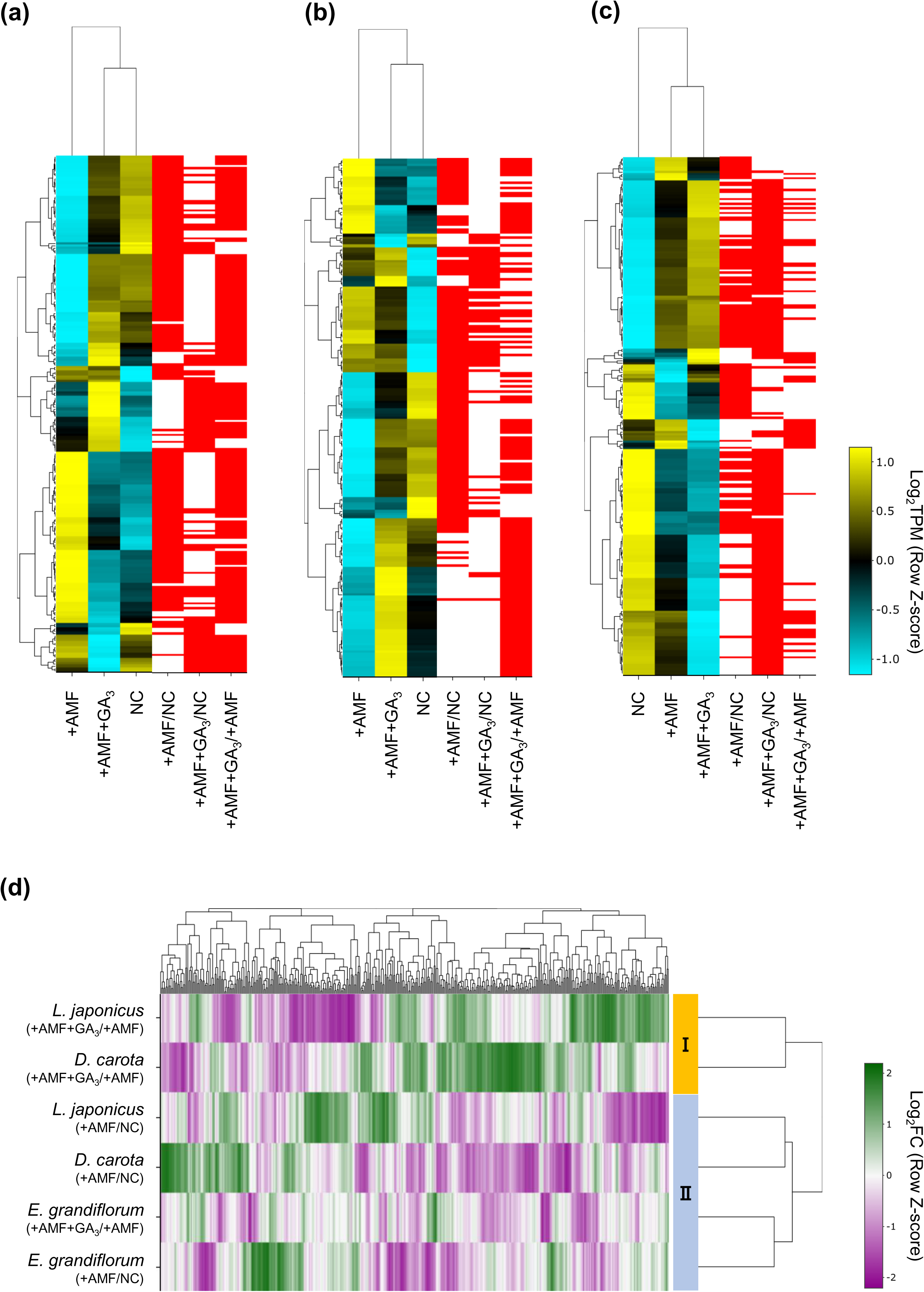
Transcriptional responses of ortholog genes to AM fungal colonization and GA treatment. The expression patterns of orthologous genes determined as DEGs in at least one condition are compared among the hosts at 6 wpi. (a–c) Individual hierarchical clustering of the mean Log_2_-transformed TPM in *L. japonicus* (a), *D. carota* (b), and *E. grandiflorum* (c). Blue, low expression level; black, mean; yellow, high expression level. The red marks in the right heatmaps represent DEGs (|Log_2_FC| > 1, FDR < 0.01). (d) Hierarchical clustering of the Log_2_FC of the total orthologs (467 genes) compared with the controls. Magenta indicates negative values, green represents positive values, and white means zero. NC, axenic conditions; +AMF, *R. irregularis* inoculation, +AMF+GA_3_, simultaneous application of *R. irregularis* inoculation and 1 µM GA_3_. The DEGs were identified by comparing +AMF with NC (+AMF/NC), +AMF+GA_3_ against NC or +AMF (+AMF+GA_3_/NC, +AMF+GA_3_/+AMF). For the TPM, Log_2_FC, and FDR values of the selected genes, see Table S5.

Given that GA-treated *E. grandiflorum* AM roots exhibited different transcriptional changes from others, we further investigated how the three plants respond to exogenous GA. In GA-treated plants, the shoots or leaves were significantly elongated, as other studies showed (Fig. S2) (Achard & Genschik, 2009; Xu *et al*., 2016; Sprangers *et al*., 2020). In addition to promoting plant growth, GA treatment has been reported to induce transcriptional changes (Cheng *et al*., 2015). Therefore, GA would have common effects on plant physiological aspects except for regulating AM symbioses.

### Comparative gene ontology enrichment analysis among examined plant species

Given that the similarity and difference in the regulation of AM symbiosis were found, we next compared the physiological functions altered in the AM roots of each host plant. Before that, DEGs in every host plant were determined by comparing +AMF with NC and +AMF+GA_3_ with +AMF conditions. The analysis showed that the number of DEGs was 4056, 3537, and 6439 in *L. japonicus*, *D. carota*, and *E. grandiflorum*, respectively (Fig. 5a). The relatively large number of DEGs in *E. grandiflorum* might be attributable to the redundant or alternative transcripts in *de novo* assembly (Duan *et al*., 2012; Ono *et al*., 2015), although the redundant contigs were removed from *de novo* reference data using CD-HIT (Li & Godzik, 2006; Tominaga *et al*., 2020b). Additionally, we classified the DEGs into three groups: A for AM fungus-responsive but GA-repressed DEGs, B for AM fungus- and GA-responsive DEGs, and C for only AM fungus-responsive DEGs (Fig. 5a). Interestingly, we found that the ratio of DEGs in Group A was relatively low in *E. grandiflorum* (3.6%) than in *L. japonicus* (25.2%) and *D. carota* (15.2%) (Fig. 5a). In contrast, the percentage of DEGs in Group B was much lower in *L. japonicus*, with 0.074%, than in *D. carota* and *E. grandiflorum*, with 2.4% and 9.0%, respectively (Fig. 5a).

**Fig. 5.**
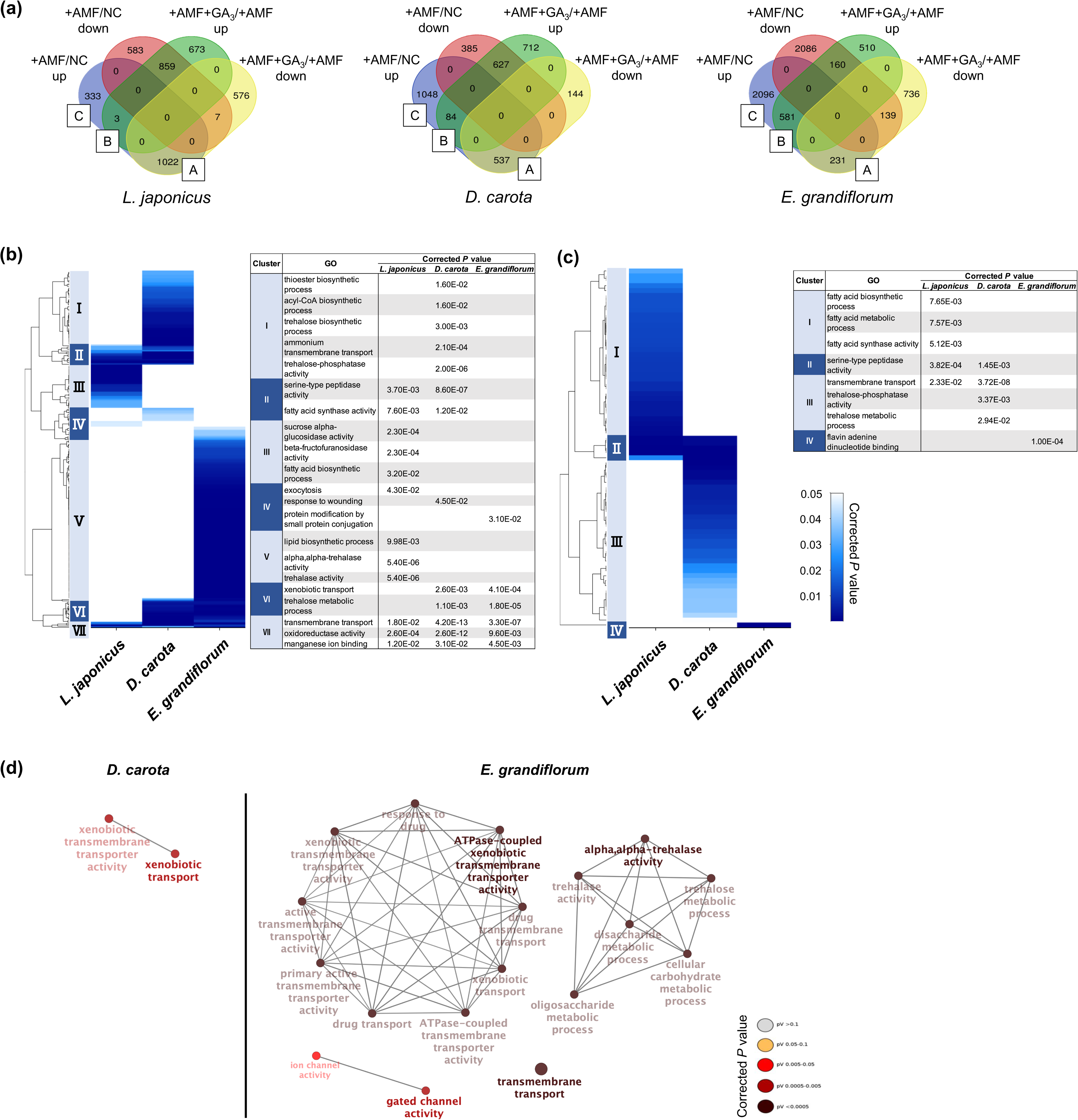
Comparisons of enriched gene ontology among AM roots of examined host plants. (a) Total DEGs (|Log_2_FC| > 1, FDR < 0.01) in individual host plants during AM symbiosis at 6 wpi. Their expression patterns classify the DEGs. The values represent the number of DEGs. As for *E. grandiflorum*, the values indicate the number of transcripts in *de novo* assembly data. NC, non-colonized control; +AMF, *R. irregularis* inoculation; +AMF+GA_3_, simultaneous application of *R. irregularis* inoculation and 1 µM GA_3_. For the determination of DEGs, transcriptomes in the host plants were compared as following: +AMF *vs.* NC (+AMF/NC), +AMF+GA_3_ *vs.* +AMF (+AMF+GA_3_/+AMF). (b, c) Hierarchical clustering of significantly enriched GO terms in the DEGs within Group A+B+C (upregulated upon AM fungal colonization) (b) and Group A (AM fungus-responsive DEGs downregulated by GA treatment) (c) in each host plant at 6 wpi (corrected *P* < 0.05). The color bar shows color-coded corrected *P*-value. (d) Categorization of GO terms in Group B (upregulated by AM fungal colonization and GA treatment) in *D. carota* and *E. grandiflorum*. No *L. japonicus* GO term was detected in the group because of the small number of DEGs. The color of circles indicates the significance of gene expression in the respective GO terms. *P*-values were calculated using a two-sided hypergeometric test in the Cytoscape plugin, ClueGO, and corrected using the Benjamini–Hochberg method. For the detailed lists of DEGs and complete GO terms in each cluster, see Table S6.

As for DEGs upregulated by AM fungal colonization (Group A+B+C), GO terms associated with membrane transport were enriched in all colonized plants (Cluster VII in Fig. 5b; Table S6). Additionally, the analysis also uncovered that peptidase- and fatty acid-related terms were also detected in *L. japonicus* and *D. carota* (Cluster II in Fig. 5b). Especially for fatty acid biosynthesis, the results are consistent with the expression pattern of *FatM* and *RAM2* in the two hosts (Fig. 2b). However, many GO terms were specific to each plant. For example, the activities of *α*-glucosidase and *β*-fructofuranosidase were shown to be upregulated upon AM fungal colonization in *L. japonicus*, indicating that the hydrolysis of disaccharides, such as sucrose, might be activated during AM symbiosis (Cluster III in Fig. 5b; Table S6). Alternatively, *D. carota* and *E. grandiflorum* promoted the trehalose biosynthetic process and trehalase activity upon fungal inoculation, respectively (Cluster I and V in Fig. 5b; Table S6). Additionally, a GO term corresponding to trehalose metabolism was found in Cluster VI. These results suggest that AM fungal colonization might influence different sugar metabolism between *L. japonicus* and the other two plants.

Furthermore, we focused on Group As to identify the functions that were upregulated upon AM fungal colonization but suppressed by exogenous GA. As illustrated in the heatmap, the DEGs in the Group A of *E. grandiflorum* were significantly enriched in GO term representing flavin adenine dinucleotide binding (GO:0050660) (Fig. 5c; Table S6). In contrast, GO terms representing transmembrane transport and peptidase were shared in *L. japonicus* and *D. carota* (Cluster II in Fig. 5c; Table S6). GO enrichment analysis again revealed that fatty acid biosynthesis was attenuated in GA-treated *L. japonicus* AM roots, corresponding to the expression pattern (Fig. 2b, 5b; Tables S4, S6). Moreover, trehalose-related processes were shown to be downregulated in GA-treated *D. carota* AM roots (Cluster III in Fig. 5c; Table S6).

Based on the positive effects of GA on AM fungal colonization and the expression levels of AM markers in *E. grandiflorum*, enriched GO terms in the Group Bs were compared with each examined plant. However, the analysis in Group B of *L. japonicus* could not be conducted because of the relatively few DEGs within the group (Fig. 5a). As illustrated in Fig. 5d, the metabolic processes of disaccharides, such as trehalose, remained upregulated in GA-treated *E. grandiflorum* AM roots. Moreover, GO terms for xenobiotic transport were enriched and shared in the Group Bs of *D. carota* and *E. grandiflorum* (Fig. 5d).

### Comparison of fungal transcriptome obtained from three examined host plants

Since a part of extraradical and intraradical hyphae were carried to RNA-seq, the resulting raw reads also contained AM fungal sequences of up to 13.7% of the total reads (Table S1). Therefore, we compared the transcriptomes of *R. irregularis* colonizing each of GA-treated *L. japonicus*, *D. carota*, and *E. grandiflorum* against one infecting the control plants. Consequently, the number of the upregulated DEGs of *R. irregularis* was relatively smaller in GA-treated *L. japonicus* than in other plants (Fig. S3a). Additionally, 8.9% of the DEGs promoted in *R. irregularis* that colonized GA-treated *E. grandiflorum* were shared with *D. carota*, whereas 4.2% of the DEGs belonged to *L. japonicus* (Fig. S3a). However, the downregulated DEGs of *R. irregularis* in GA-treated *L. japonicus* were mostly shared with *D. carota* (24.7%) compared with *E. grandiflorum* (9.9%) (Fig. S3a).

*Rhizophagus irregularis* seemed to differentially respond to the examined plants; therefore, we next conducted GO enrichment analysis on the fungal DEGs. Hierarchical clustering arranged by Log_2_FC showed six clusters depending on the expression patterns of DEGs (Fig. S3b). GO terms associated with a mitogen-activated protein kinase (MAPK) activity were often detected in Cluster IV, where upregulated DEGs in *R. irregularis* associating with GA-treated *E. grandiflorum* were dominant (Fig. S3b; Table S7). Additionally, glycogen metabolism- and wax biosynthesis-related terms were found in the cluster (Fig. S3b; Table S7). Interestingly, DEGs in Cluster VI, where numerous downregulated DEGs were found in GA-treated *L. japonicus*, were enriched in some GO terms corresponding to the elongation and oxidation of fatty acid (Fig. S3b; Table S7). This indicates that the allocation of host-derived fatty acids would be attenuated by GA application in *L. japonicus*, which could be explained by GA-suppressed fatty acid biosynthesis in the plant AM roots (Fig. 3b; Table S4).

## Discussion

In this study, we performed comparative transcriptomics and revealed that the transcriptional response during AM symbiosis was similar among *L. japonicus*, *D. carota*, and *E. grandiflorum* AM roots in the control condition. Moreover, a certain set of known genes conserved in every tested plant were transcriptionally promoted upon AM fungal colonization; this is similar to previous comparative transcriptomics between several AM hosts (Fig. 2) (Sugimura & Saito, 2017; An *et al*., 2018). Another study revealed the conservation of *RAD1*, *STR*, and *STR2* in AM host lineages and suggested their common functions in AM symbiosis; the AM symbiosis-related genes conserved in the examined plants would comparably function in establishing AM symbiosis as well (Radhakrishnan *et al*., 2020). Additionally, most selected TFs showed significantly upregulated expression levels upon AM fungal inoculation in the examined plants (Fig. 2b–d). *CBX1*, *WRI5s*, *RAM1*, and *RAD1* have been known to be involved in the full development of arbuscule by upregulating the expression of downstream genes, such as *PT4*, *RAM2*, and *STR*, in legume and monocot model plants (Gobbato *et al*., 2012; Park *et al*., 2015; Rich *et al*., 2015; Pimprikar *et al*., 2016; Luginbuehl *et al*., 2017; Rich *et al*., 2017; Jiang, Y *et al*., 2018; Xue *et al*., 2018). Therefore, the TFs conserved in *D. carota* and *E. grandiflorum* might function in accommodating AM fungi and arbuscule formation and the model plants. In addition to transcriptional regulation, as previously shown, growth promotion was observed regardless of the distinct AM morphotypes (Fig. 1i) (Hong *et al*., 2012). This result would be supported by the fact that phosphate and ammonium transporters were highly expressed in each host plant colonized by *R. irregularis* (Fig. 2b–d). Taken together, our study provided knowledge supporting that the selected genes would act to establish AM symbiosis in the phylogenetically distant host plants with various AM morphotypes.

Nevertheless, this study clarified that GA treatment oppositely modulated the AM fungus-responsive transcriptional changes between *L. japonicus*/*D. carota* and *E. grandiflorum*, which are reminiscent of the colonization traits (Figs. 1g, 2b, 4). Recently, CYCLOPS required for both AM symbiosis and root nodule symbiosis has been reported to bind the *cis*-element on *LjRAM1* promoter and upregulate gene expression in concert with a Ca^+2^/calmodulin-dependent protein kinase (CCaMK) and a GA-degradable repressor of GA signaling, DELLA protein (Silverstone *et al*., 2001; Achard & Genschik, 2009; Pimprikar *et al*., 2016). The involvement of DELLA in the complex is thought to trigger the GA-mediated inhibition of *RAM1* expression, resulting in the severely suppressed AM fungal accommodation. *Daucus carota* showed reduced rates of AM fungal colonization and expression levels of *DcRAM1* and *L. japonicus*, indicating that the transcriptional regulation of downstream genes would be common (Fig. 6). However, the expression levels of *RAM1* and some of the downstream genes were significantly or slightly promoted in GA-treated *E. grandiflorum* (Fig. 2d). This implies that DELLA protein might not be necessary or inhibit the expression of downstream genes in this plant. Moreover, other TF genes, such as *CBX1* or *RAD1*, might upregulate the downstream genes instead of RAM1 in *E. grandiflorum*. Primarily, *RAD1* would act as an alternative to *RAM1* because some AM host plants, such as *Marchantia paleacea*, lack *RAM1* but possess *RAD1* in their genomes (Grosche *et al*., 2018; Radhakrishnan *et al*., 2020). To clarify the upstream regulation of these TFs in *E. grandiflorum*, further investigation would be necessary.

**Fig. 6.**
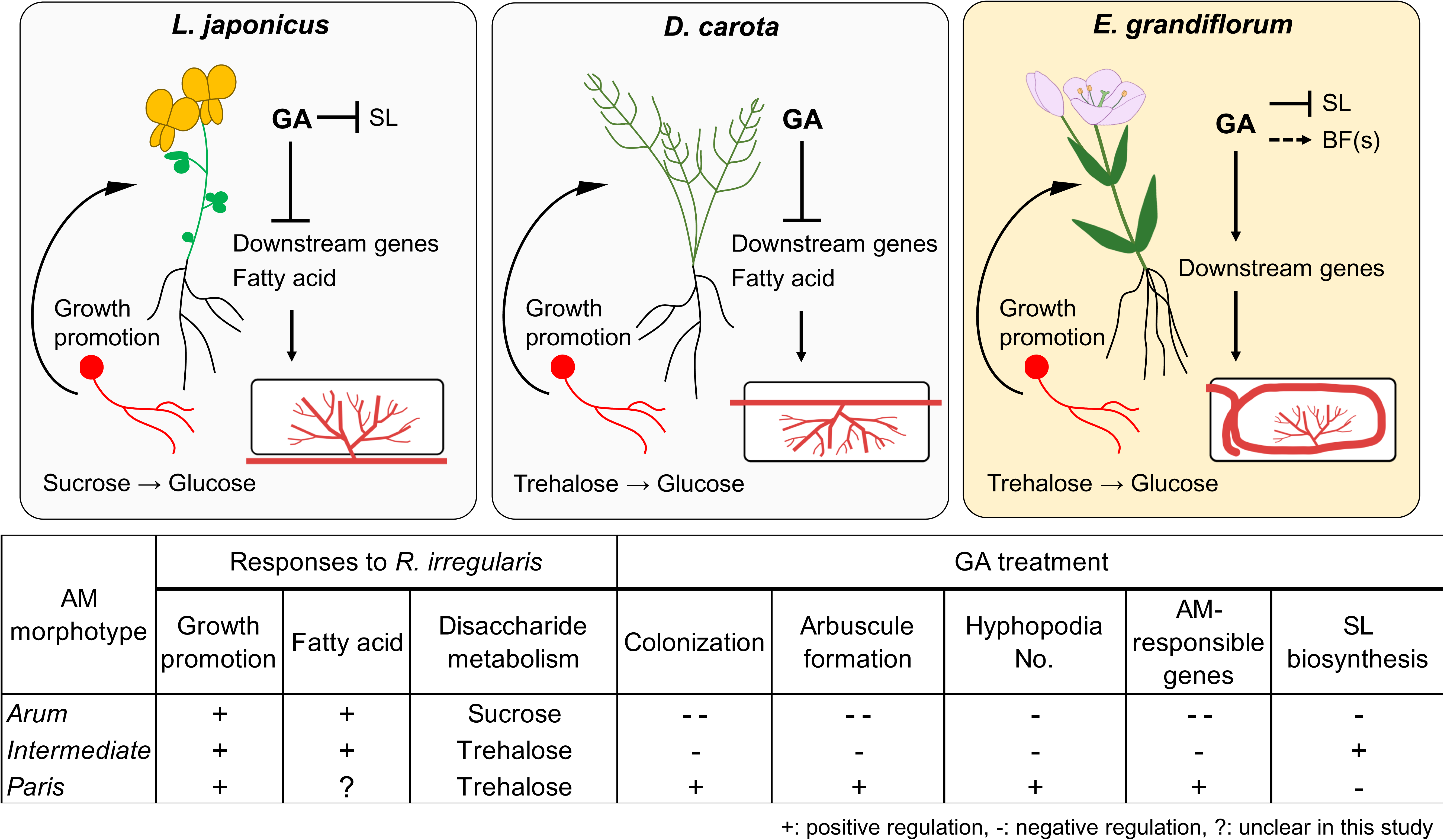
Proposed model for conserved and divergent responses to AM fungi and exogenous GA in genetically distant host plants at 6 wpi. The hypothetical model and table represent common and different responses to *R. irregularis* colonization and exogenous GA treatment in *L. japonicus* (*Arum*-type), *D. carota* (Intermediate-type), and *E. grandiflorum* (*Paris*-type). AM fungal colonization contributed to growth promotion in each tested plant forming the distinct AM morphotypes. However, GA treatment suppressed but enhanced a set of conserved genes regulating AM symbiosis in *L. japonicus*/*D. carota* and *E. grandiflorum*, respectively. On the other hand, GA transcriptionally inhibited SL biosynthetic genes in *L. japonicus* and *E. grandiflorum*, implying unidentified branching factor in *E. grandiflorum*. Moreover, disaccharide broken down during AM symbiosis might be different: sucrose and trehalose in *L. japonicus* and *D. carota*/*E. grandiflorum*. +, positive regulation; -, negative regulation; ?, response of the tested plants that could not be cleared in this study.

AM host plants exudate some signal molecules to the rhizosphere to enhance the association with symbiotic fungi and other microbes, resulting in growth promotion in the hosts (Lagrange *et al*., 2001; Akiyama *et al*., 2005; Nakayasu *et al*., 2021). The representative host-derived signal molecules toward AM fungi, SL, are required for the formation of AM fungal hyphopodia, and its biosynthesis and exudation are inhibited by exogenous GA (Ito *et al*., 2017; Kobae *et al*., 2018). This study indicated that GA treatment would enhance SL production in *D. carota* despite the fact that the number of invading hyphopodia was reduced in GA-treated *D. carota* (Figs. 1h, 2c, S1). Since SL biosynthesis has a negative feedback regulation, GA-repressed SL production might trigger the increase in *DcCCD8* expression upon exogenous GA at 6 wpi (Hayward *et al*., 2009; Proust *et al*., 2011). Alternatively, the reduction of SL exudation by GA might have occurred at an earlier time point than 6 wpi in *D. carota*. However, we could confirm the inhibitory effects of GA on SL biosynthetic genes in *L. japonicus* and *E. grandiflorum* (Figs. 3b–d, S1). Nevertheless, the number of the hyphopodia was drastically induced in GA-treated *E. grandiflorum* roots, implying the existence of yet-to-be-identified signal molecule(s) except for SL (Fig. 1h) (Tominaga *et al*., 2020a). We could not identify the unknown signal molecule(s); however, the possible existence was assumed from the GO enrichment analysis on *R. irregularis*. As shown in Fig. S3b, some GO terms representing the activity of MAPK kinase kinase, MAPK Kinase, and MAPK were detected in *R. irregularis* colonizing GA-treated *E. grandiflorum* (Table S7). In plant pathogenic fungi, such as *Ustilago maydis* and *Magnaporthe oryzae*, MAPK cascade is required for the formation of appressoria and their virulence after perceiving some signal molecules derived from the host plants (Hamel *et al*., 2012; Li *et al*., 2017; Jiang *et al*., 2018). Although the necessity of the MAPK cascade in *R. irregularis* hyphopodia formation remains unclear, the fungus might sense some signal molecule(s) different from SLs exudated from GA-treated *E. grandiflorum* roots.

In nature, it has been known that host plants forming *Arum*-type AM are often seen in vacant areas, whereas those showing *Paris*-type AM prefer relatively dark space, such as forest floor (Yamato & Iwasaki, 2002; Yamato, 2004). Interestingly, the relationship between light condition and symbiotic interactions has been indicated. For instance, AM fungal colonization and SL production are attenuated by far-red or *phyB* mutation in *L. japonicus* and tomato (Nagata *et al*., 2015). Moreover, shaded areas make plants accumulate GA to elongate their stems for the light (Hisamatsu *et al*., 2005; Bou-Torrent *et al*., 2014; Colebrook *et al*., 2014; Li *et al*., 2017; Yang & Li, 2017). Taken together, the unique regulatory mechanisms and unidentified signal molecule(s) in *E. grandiflorum* might be exploited to adapt to the conditions of their habitats. However, *Cardamine hirsute* adapted to shaded area possesses a mechanism that attenuates shade-induced hypocotyl elongation (Paulišić *et al*., 2021). Thus, the relationship between GA-promoted *Paris*-type AM symbiosis and environmental cues should be investigated.

The loss of genes encoding enzymes required for polysaccharide degradation in AM fungi demands host plants on glucose (Kobayashi *et al*., 2018). In arbuscules containing cells of *M. truncatula*, AM fungi-responsive localization and the expression of *MtSucS1* and *MtSWEET1b* are thought to produce glucose and export it toward AM fungi between the symbiotic interface (An *et al*., 2019). In fact, our GO enrichment analysis showed activated sucrose hydrolysis during AM symbiosis in *L. japonicus*, which was supported by AM-induced *LjSucS1* and *LjSWEET1b*s expression (Fig. 2b; Table S4). Alternatively, another disaccharide, trehalose, was indicated to be broken down in *D. carota* and *E. grandiflorum* during AM symbiosis as the AM-induced activities and expressions of plant trehalase (*TRE1*) were seen in the host plants (Fig. 5b, d; Tables S4, S6). This difference might be attributable to the intracellular hyphal invasion in Intermediate- and *Paris*-type AM roots (Fig. 1c–f). However, the increase in *TRE1* expression was also observed in *L. japonicus* AM roots at 6 wpi (Table S4). Interestingly, it is known that most of the storage carbohydrates found in fungi are trehalose, and AM fungi can synthesize and accumulate trehalose in the spores and hyphae (Shachar-Hill *et al*., 1995; Bago *et al*., 1999; Pfeffer *et al*., 1999; Kameoka *et al*., 2019). Additionally, the upregulation of *TRE1* expression has also been seen in *Arabidopsis thaliana* infected by a pathogenic fungus, *Plasmodiophora brassicae*, which is considered as the maintenance of sugar concentration and physiological homeostasis in the roots by removing fungal-derived trehalose (Brodmann *et al*., 2002). Although we cannot exclude that trehalose is used as a carbon resource for AM fungi, TRE1 might be required to reduce AM fungi-derived trehalose concentration in the host plants.

In summary, a particular set of conserved AM symbiosis-related genes would commonly function to accommodate AM fungi in the phylogenetically distant AM host plants, thereby forming distinct AM morphotypes. However, our study uncovered that GA oppositely affects the transcriptions of the conserved AM symbiosis-related genes: negative in *Arum*- and Intermediate-AM symbioses in *L. japonicus* and *D. carota*, respectively, and positive in *Paris*-type AM symbiosis in *E. grandiflorum* (Fig. 6). These findings advance the comprehensive understanding of transcriptomic regulation and the diversity of GA-mediated effects on AM symbioses among host plants.

## Acknowledgment

We would like to thank Dr. Gabriela Bindea (Cordeliers Research Center) for preparing the list and enabling us to conduct GO enrichment analysis on *L. japonicus* and *E. grandiflorum*. Additionally, we appreciate the National BioResource Project (Legume Base) and Dr. Satoko Yoshida (Nara Institute of Science and Technology) for kindly providing *L. japonicus* and *O. minor* seeds, respectively. This work was partially supported by the NIBB Cooperative Research Programs (Next-generation DNA Sequencing Initiative: 19-433, 20-407) and JSPS KAKENHI Grant-in-Aid for JSPS Fellows (Grant No. 20J21994 to TT).

## Author Contribution

TT and HK designed the experiments; TT, YS, YH, and MA performed the experiments; TT, CM, KY, and SS analyzed the sequencing data; TT, CM, MA and HK wrote the manuscript. All authors approved the final manuscript.

## Data availability

The nucleotide sequence data obtained from our transcriptome analysis has been deposited into the DDBJ Sequence Read Archive under the accession number DRA012117. *De novo* assembly and annotation list of *E. grandiflorum* are available on Open Science Foundation with DOI 10.17605/OSF.IO/TQ7XJ or https://osf.io/tq7xj/?view_only=b6bec888fd80417ea636c3b6b58f07c1.

## Supporting Information

**Fig. S1** Germination rate of *O. minor* treated with the root extracts of the examined host plants.

**Fig. S2** Phenotyping of GA-treated host plants.

**Fig. S3** Transcriptional responses to GA-treated host plants in *R. irregularis*.

**Table S1** Results of trimming and mapping of RNA-seq read.

**Table S2** Lists of SonicParanoid outputs.

**Table S3** List of BLASTp results and selected genes in this study.

**Table S4** Expression patterns of the selected genes in the examined host plants.

**Table S5** Expression levels and fold changes of orthologs in Fig. 4.

**Table S6** Enrichment analysis of the examined host plants colonized by *R. irregularis* and treated with GA.

**Table S7** Transcriptomic analysis of *R. irregularis* colonizing examined host plants in the presence of GA.

