## Supplemental Figure 1-3 for "Conservation and diversity in transcriptional responses among host plants forming distinct arbuscular mycorrhizal morphotypes"

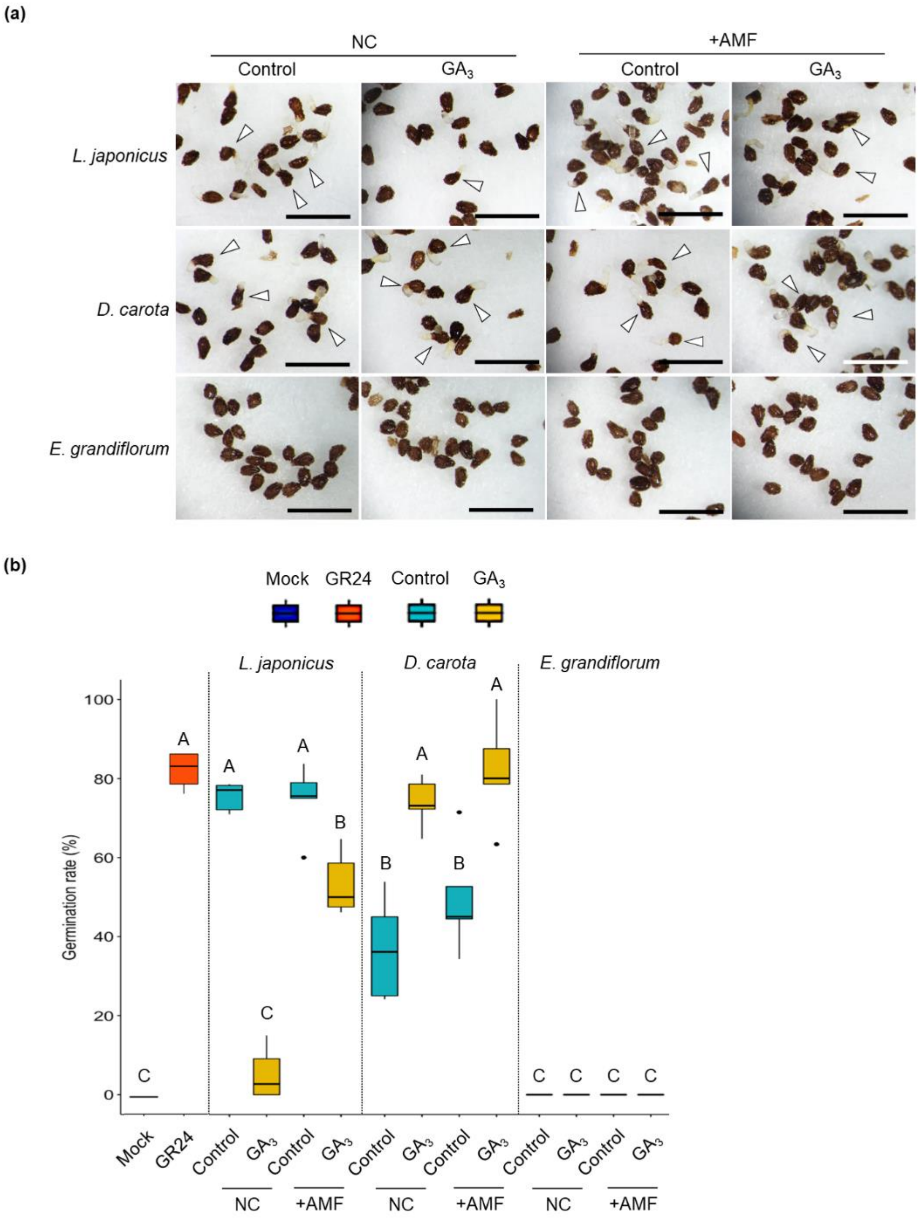


**Fig. S1 Germination rate of *O. minor* treated with the root extracts of the examined host plants.** The root extracts from 6-wk-old *L. japonicus*, *D. carota*, and *E. grandiflorum* roots were treated to *O. minor* seeds for 5 d. The host plants were grown under axenic (NC) and monoxenic conditions with *R. irregularis* (+AMF) in the absence and presence of 1 µM $GA_3$. Acetone (mock) and 1 µM *rac*-GR24 were applied for negative and positive controls, respectively. (a) Images of *O. minor* seeds treated with each sample. Arrowheads indicate germinating *O. minor* seeds. Scale bars, 1 mm. (b) Germination rate (%) of *O. minor* treated with each sample. The different alphabets represent significant differences at $P < 0.05$ in the Tukey–Kramer test (n = 5).

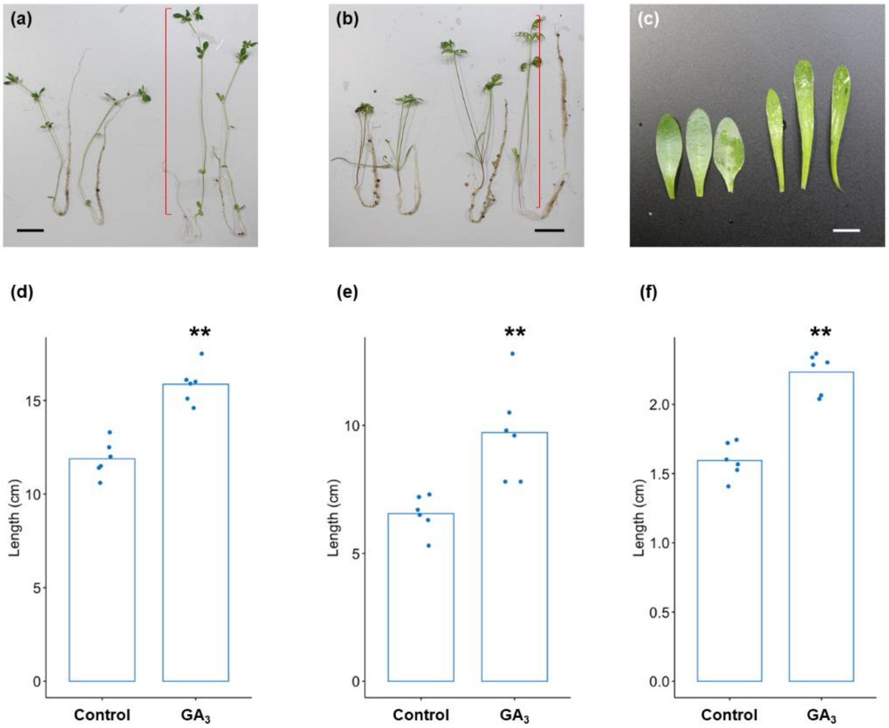


**Fig. S2 Phenotyping of GA-treated host plants.** The length of *L. japonicus* and *D. carota* shoots or *E. grandiflorum* leaves were measured. These plants were treated with 0.01% ethanol as for the control and 1 µM $GA_3$ and grown for 6 wk. (a–c) Images of GA-treated seedlings of *L. japonicus* (a), *D. carota* (b), or third leaves of *E. grandiflorum* (c). Bars, 2 cm (a, b), 5 mm (c). (d–f) The length of *L. japonicus* (d) and *D. carota* (e) shoot or of *E. grandiflorum* leaf (f). Bars and plots indicate the mean and individual values, respectively. Asterisks show significant differences in Wilcoxon rank-sum test (**: $P < 0.01$).

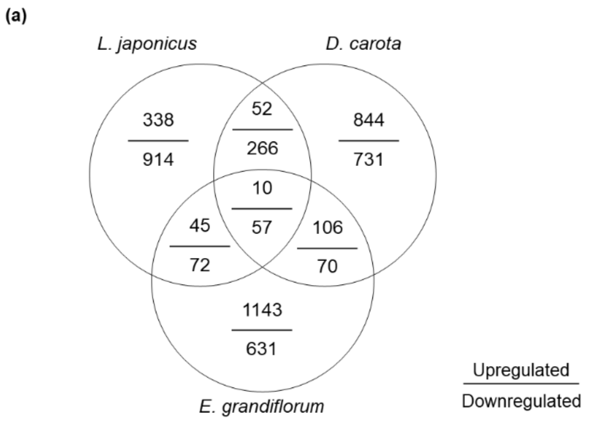


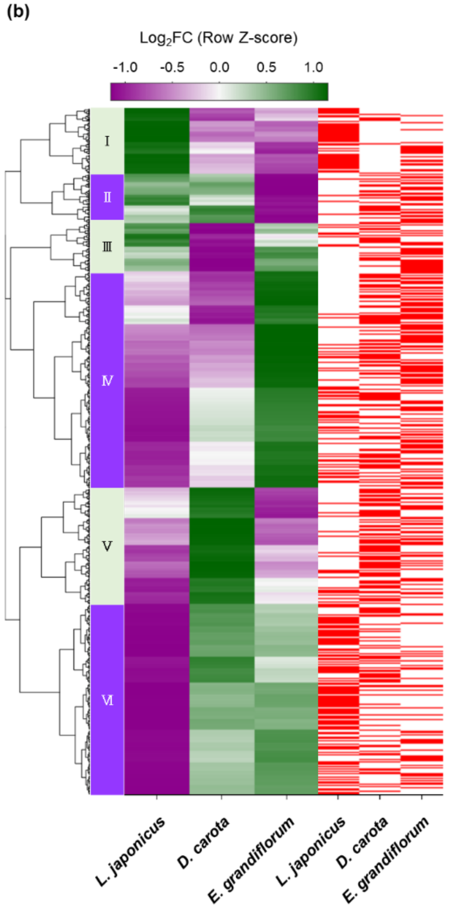


| Cluster | GO | *P* value |
|---|---|---|
| I | glycogen metabolic process | 0.000940 |
| | lipid storage | 0.00398 |
| II | carotenoid biosynthetic process | 0.00101 |
| | aldehyde catabolic process | 0.00167 |
| III | chitin catabolic process | 0.00771 |
| | chitosan layer of spore wall | 0.00141 |
| | chitin binding | 0.00185 |
| | chitin deacetylase activity | 0.00214 |
| IV | protein phosphorylation | 2.10E-08 |
| | wax biosynthetic process | 0.00156 |
| | cell-cell adhesion | 0.00221 |
| | interaction with host | 0.00258 |
| | activation of MAPKK activity | 0.00274 |
| | response to xenobiotic stimulus | 0.00507 |
| | glycogen metabolic process | 0.00537 |
| | protein secretion | 0.00568 |
| | glycerol-3-phosphate dehydrogenase complex | 0.00873 |
| | MAP kinase kinase activity | 0.001200 |
| | MAP kinase kinase kinase activity | 0.00138 |
| | lipase activity | 0.00451 |
| V | chitin metabolic process | 0.006900 |
| | chitin synthase activity | 2.70E-05 |
| | chitin binding | 0.000320 |
| VI | fatty acid omega-oxidation | 5.80E-09 |
| | fatty acid elongation | 0.000170 |
| | pantothenate biosynthetic process | 0.00147 |
| | fatty acid beta-oxidation | 0.00308 |
| | lipid droplet | 3.20E-06 |

**Fig. S3Transcriptional responses to GA-treated host plants in *R. irregularis*.** The *R. irregularis*-derived reads obtained from total reads of the host plants were subjected to transcriptomic analysis. The ratio of AM fungal reads is summarized in Table S1. (a) Identification of DEGs ($|Log_2FC| > 1$, FDR < 0.05) in *R. irregularis* colonizing 1 µM $GA_3$-treated *L. japonicus*, *D. carota*, and *E. grandiflorum* compared when it infected the 0.01% ethanol-treated (control) plants at 6 wpi. The values represent the number of DEGs upregulated or downregulated by GA treatment. (b) Heatmap showing the hierarchical clustering of AM fungal genes that significantly expressed in at least one treatment compared with the control conditions (4764 genes). The Z-score-normalized $Log_2FC$ are arranged by their expression patterns. Magenta indicates negative values, green represents positive values, and white means zero. The right tables show GO terms that significantly enriched in Clusters IV and VI. *P*-values were calculated using the Kolmogorov–Smirnov test, and significant GO terms were identified using the *Elim* method.
